# *Deformed wing virus* type A, a major honey bee pathogen, is vectored by the mite *Varroa destructor* in a non-propagative manner

**DOI:** 10.1101/660985

**Authors:** Francisco Posada-Florez, Anna K. Childers, Matthew C. Heerman, Noble I. Egekwu, Steven C. Cook, Yanping Chen, Jay D. Evans, Eugene V. Ryabov

**Author notes:** **Corresponding authors:** Francisco Posada-Florez, USDA, Agricultural Research Service, Bee Research Lab, BARC-East Bldg. 306, 10300 Baltimore Ave., Beltsville, MD 20705, USA, Eugene V. Ryabov, USDA, Agricultural Research Service, Bee Research Lab, BARC-East Bldg. 306, 10300 Baltimore Ave., Beltsville, MD 20705, USA.

## Abstract

Honey bees, the primary managed insect pollinator, suffer considerable losses due to *Deformed wing virus* (DWV), an RNA virus vectored by the mite *Varroa destructor*. Mite vectoring has resulted in the emergence of virulent DWV variants. The basis for such changes in DWV is poorly understood. Most importantly, it remains unclear whether replication of DWV occurs in the mite. In this study, we exposed *Varroa* mites to DWV type A via feeding on artificially infected honey bees. A significant, 357-fold increase in DWV load was observed in these mites after 2 days. However, after 8 additional days of passage on honey bee pupae with low viral loads, the DWV load dropped by 29-fold. This decrease significantly reduced the mites’ ability to transmit DWV to honey bees. Notably, negative-strand DWV RNA, which could indicate viral replication, was detected only in mites collected from pupae with high DWV levels but not in the passaged mites. We also found that *Varroa* mites contain honey bee mRNAs, consistent with the acquisition of honey bee cells which would additionally contain DWV replication complexes with negative-strand DWV RNA. We propose that transmission of DWV type A by *Varroa* mites occurs in a non-propagative manner.

## INTRODUCTION

Honey bee (*Apis mellifera*), the primary managed insect pollinator, suffers considerable losses caused by pathogens^1,2^, including *Deformed wing virus* (DWV)^3^, which endanger pollination service worldwide^4,5^. Three “master” variants of DWV were discovered including DWV-A^3^, DWV-B (also known as *Varroa destructor virus*-1^6^), and DWV-C^7^. In addition, recombinants between DWV-A and DWV-B were reported as major DWV variants in some localities in the UK, Israel and France^8–10^. Negative impacts of DWV infection usually manifests when the virus is vectored by the ectoparasitic mite *Varroa destructor*^11–13^. Importantly, both DWV-A and DWV-B variants are vectored by the mite. The Varroa mite, originally a parasite of the Asian honey bee *Apis cerana*, spread to the Western honey bee *A. mellifera* in the beginning of the 20th century, reaching Western Europe and subsequently North America about 30 years ago^14^. DWV was present in honey bees prior to *Varroa* arrival, although at low levels^15^. The mite, which provides an efficient horizontal transmission route for DWV, transformed this covert virus into a highly virulent virus. DWV levels in Varroa-infested bees are 100 to 1000-fold higher compared to Varroa-free bees^16,17^. Such increases of DWV virulence were accompanied by genetic changes in the virus, as reported in Hawaii^15^ and Scotland^16^, then further confirmed by an analysis of DWV evolution worldwide^117^.

Unraveling the mechanisms that drive genetic changes in DWV requires a comprehensive understanding of the interactions between each component within the tripartite system “Honey bee-DWV-*Varroa* mite”. Importantly, there is no agreement on whether replication of all DWV variants takes place within the mite vector^17^. The currently accepted view that DWV replication occurs in *Varro*a mites is largely based mostly on the detection of negative-strand DWV RNA^16,18,19^. DWV belongs to the genus *Iflaviridae*, order *Picornavirales*, members of which are known to have virus particles containing exclusively positive strands as their genomic RNA^20,21^. Therefore, presence of negative-strand viral RNA in the mites could indicate that replication of DWV takes place in this species. Virus particles were reported in the cells of Varroa mites possibly harboring DWV-B (VDV1)^6^. Contrary to this, an immunolocalization study of *Varroa* mites feeding on DWV-infected pupae revealed no presence of DWV in *Varroa* cells. Instead, DWV antigen was detected exclusively in the mites’ midgut lumen^22^. This apparent disagreement could be a explained by the use of different DWV isolates (i.e. “master types”), which could differ in respect to their replication in Varroa mites. For example, recently negative strand RNA of DWV-B was detected in*Varroa* synganglions^23^. Unfortunately, previous studies which investigated DWV replication in the mites, did not include identification of the DWV type^18,19^. Although strong positive correlation of DWV loads was reported in naturally infested colonies between the *Varroa* mites and associated honey bee pupae, this cannot be used as an argument in support of DWV replicates within *Varroa* mainly because DWV is vectored by mites and could be acquired by the mites from the same pupae which they infect. Therefore, the question of whether replication of DWV-A, the most widespread variant, occurs in the mite vector remains open.

In this study, we investigated temporal dynamics of both positive and negative-strand DWV-A RNA accumulation in *Varroa* mites and honey bee pupae, as well as transmission of the virus between mites and pupae. The experiments were conducted in controlled conditions rather than in the honey bee colony environment, which allowed for feeding of individual mites only on certain pupae for a chosen period of time. By using artificial infection of the honey bee pupae with DWV-A, we could specifically monitor acquisition of the virus by the mites and transmission to new pupae. Moreover, we were able to trace the DWV transmission route by using a cloned DWV isolate. We found no evidence supporting DWV-A replication in the *Varroa* mites, demonstrating that virus loads in the mite are highly dynamic, and that the mites’ ability to transmit DWV significantly declines following loss of their DWV loads after feeding on the honey bees with low DWV levels.

## Results

### Significant increases and decreases in DWV loads in *Varroa* mites following feeding on pupae with different DWV levels

Two experiments were carried out to investigate the effect of pupal DWV levels on mite DWV loads (Figs. 1 and 2). For both experiments, phoretic mites were collected from adult honey bees in colonies with DWV levels typical for US honey bee colonies. A control sample of phoretic mites (V-Pho_1_) was immediately collected for molecular analysis. Two or three mites were placed on individual honey bee pupae which had been injected 3 days earlier with either DWV or a control PBS solution. The virus used was a clone-derived DWV-304(GenBank Accession Number MG831200) (E. Ryabov, personal communication), which was a strain of DWV type A^7^ isolated from a typical *Varroa*-infested honey bee colony. The mites and pupae were incubated in individual gelatin capsules as described earlier^24^ (Fig. 1B). After 2 days, all injected pupae (P0-PBS_1_, P0-DWV_1_, P0-PBS_2_ and P0-DWV_2_) and one *Varroa* mite from each pupa (V0-PBS_1_, V0-PBS_2_, V0-DWV_1_ and V0-DWV_2_) were collected. The remaining one or two mites from each injected pupa were transferred to fresh pupae for an additional 2 days of feeding. These two-day passages were repeated four times, after which V*arroa* mites (V4-PBS_1_, V4-PBS_2_, V4-DWV_1_ and V4-DWV_2_) and pupae (P4-PBS_1_ and P4-PBS_1_) were collected for molecular analysis. The mites’ survival rates in Experiments 1 and 2 were 95.7% and 83.3%, respectively; with no significant difference observed between the mites initially exposed to pupae injected with DWV or PBS (Experiment 1, p = 0.32; Experiment 2, p = 0.93; Wilcoxon test). High *Varroa* survival in these experiments demonstrates that mites were robust under *in vitro* conditions, allowing for reliable virus accumulation and transmission results.

**Figure 1.**
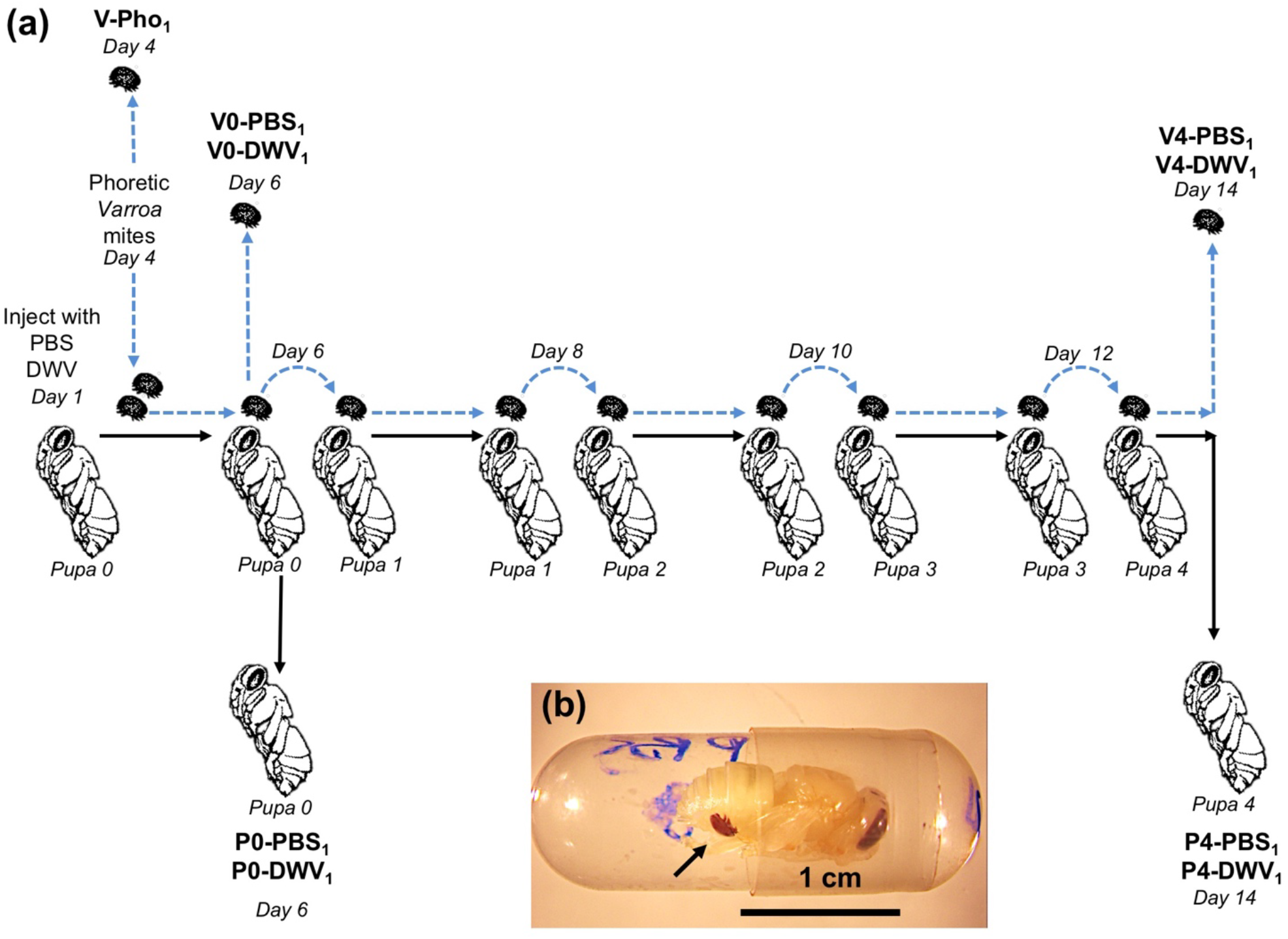
Experiment 1 design: “*Varroa* mite DWV acquisition experiment”. **(a)** Honey bee and the mite treatment groups are shown in bold. Times of sampling (days of the experiment) are indicated. **(b)** *In vitro* incubation of honey bee pupa with *Varroa* mites in gelatin capsule.

**Figure 2.**
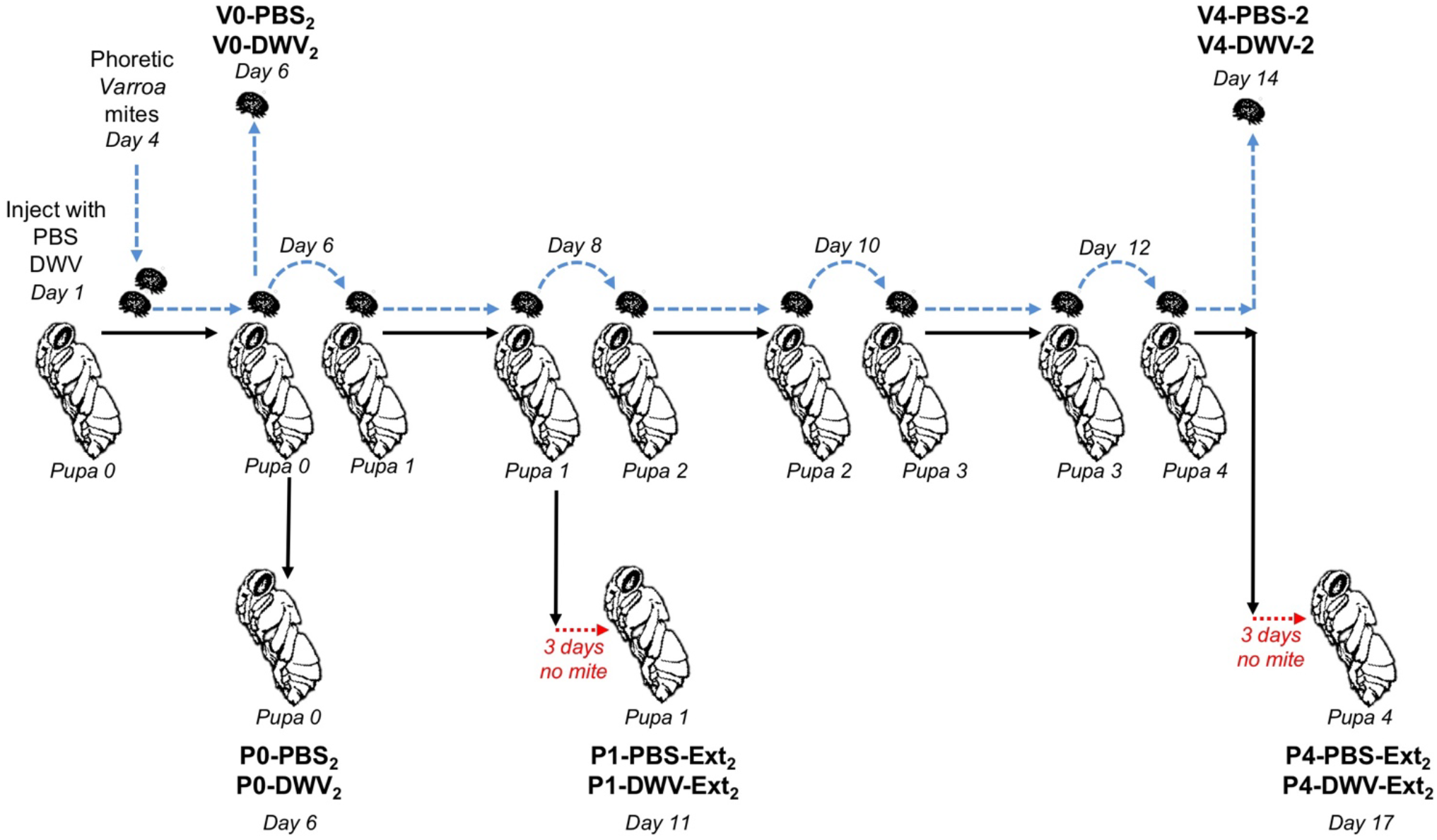
Experiment 2 design: “*Varroa-*mediated DWV transmission experiment”. Honey bee and the mite treatment groups are shown in bold. Times of sampling (days of the experiment) are indicated.

We quantified copy numbers of DWV RNA in the collected mites and pupae by using RT-qPCR assays which included reverse transcription using random primers and detected mainly positive-strand genomic RNA. The levels of DWV in pupae injected with DWV were significantly higher than in the PBS-injected pupae (Fig. 3c, Fig. 4c). In particular, we confirmed development of high-level DWV infection in the DWV-injected groups, with 19 out of 20 bees in P0-DWV_1_, and 11 out of 12 bees in P0-DWV_2_ containing more than 10^9^ DWV genomes copies. PBS-injected bees had mainly low levels of DWV with only 1 out of 20 bees in P0-PBS_1_, and 1 out of 12 bees P0-PBS_2_ having more than 10^9^ DWV genome copies. The level of 10^9^ DWV genome copies was used as the accepted threshold infection level between asymptomatic and symptomatic DWV infections (i.e. “covert” and “overt”) as reported in several studies^19,25,26^. Therefore, we obtained groups of honey bee pupae with significantly different DWV levels to investigate DWV acquisition and transmission by the mites.

**Figure 3.**
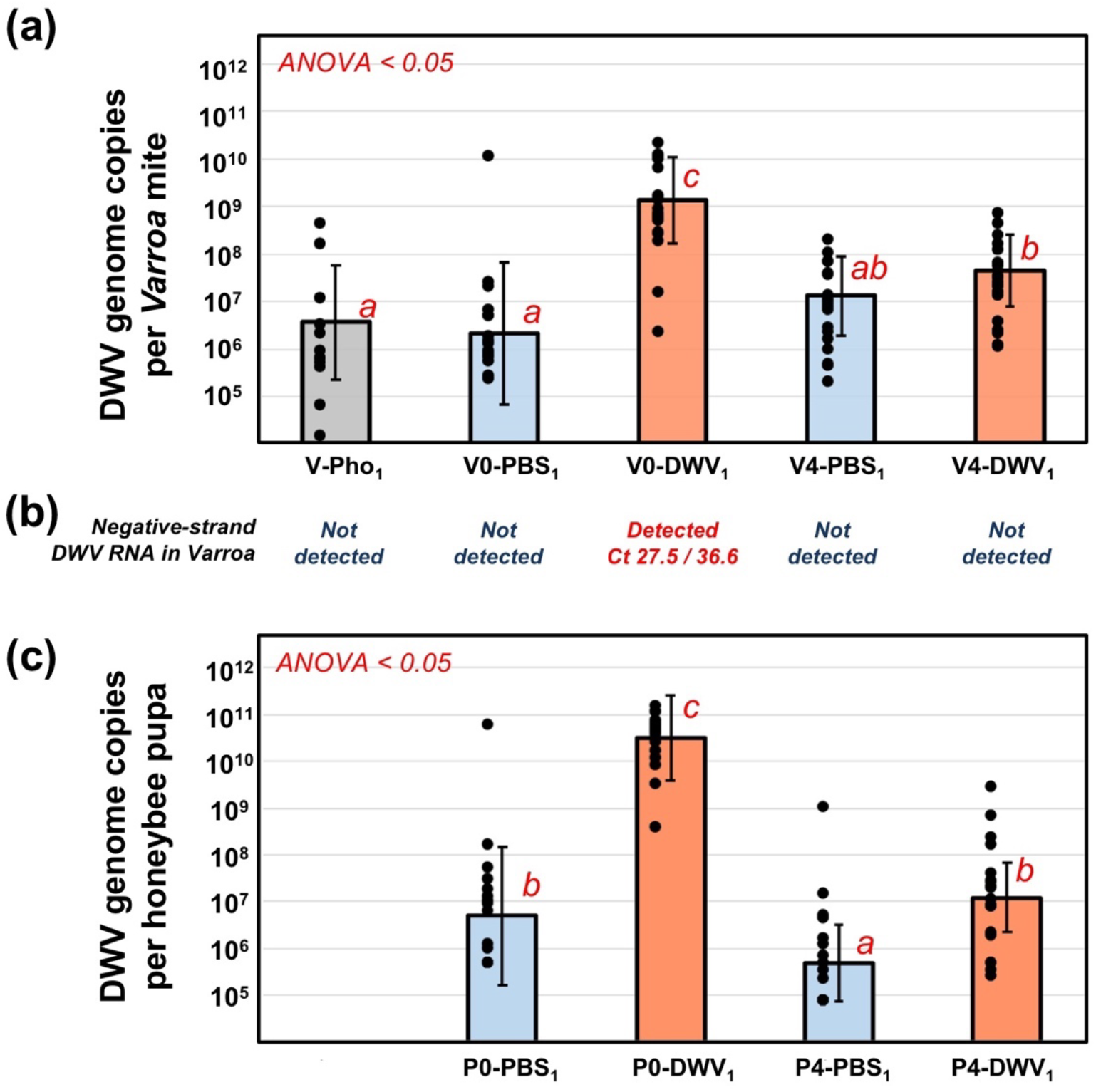
Dynamics of DWV RNA accumulation in *Varroa destructor* mites and honey bee pupae from Experiment 1: “*Varroa* mite DWV acquisition experiment”. **(a, c)** Average DWV RNA copies per **(a)** *Varroa destructor* mites, and **(b)** honey bee pupae, ±SD. Red letters above the bars indicate significantly and non-significantly different groups (ANOVA). Treatment groups are named as in the Fig. 1. Individual mite and pupa copy numbers indicated by black dots. Corresponding mite **(a)** and pupae **(c)** groups are aligned vertically. **(b)** Detection of negative strand DWV RNA in *Varroa* mite groups. The Ct values shown are as follows: Assay 1 / Assay 2.

**Figure 4.**
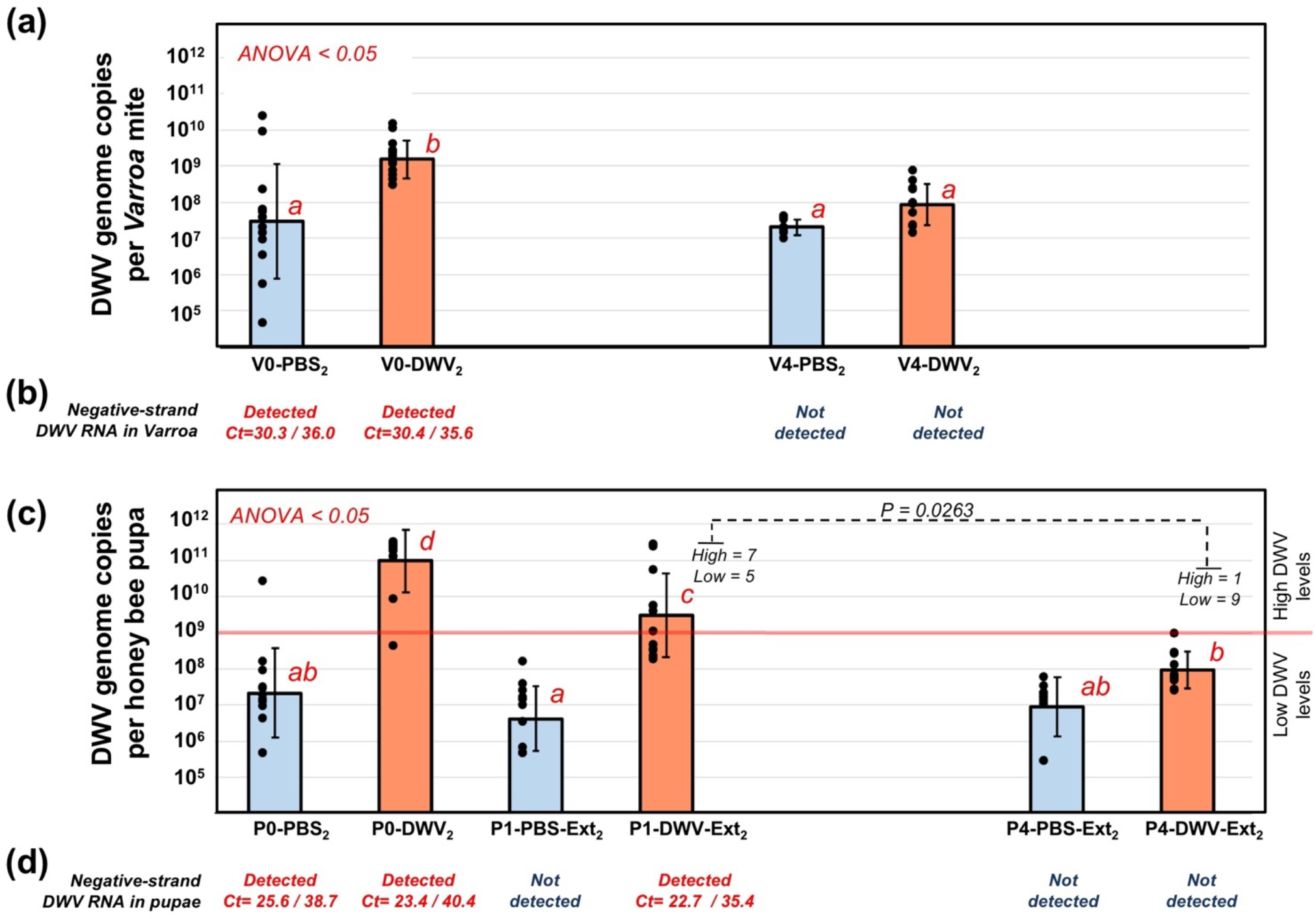
Dynamics of DWV RNA accumulation in *Varroa destructor* mites and honey bee pupae from Experiment 2: “*Varroa-*mediated DWV transmission experiment”. **(a, c)** Average DWV RNA copies per **(a)** *Varroa destructor* mites, and **(b)** honey bee pupae, ±SD. Red letters above the bars indicate significantly and non-significantly different groups (ANOVA). Treatment groups are named as in Fig. 2. Individual mite and pupa copy numbers indicated by black dots. Corresponding mite **(a)** and pupae **(c)** groups are aligned vertically. The threshold levels of 10^9^ DWV genomic RNA copies per honey bee pupae is shown as a red line in **(c).** Detection of the negative strand DWV RNA in **(b)** *Varroa* mite groups and **(d)** honey bee pupae. The Ct values shown are as follows: Assay 1/ Assay 2.

Quantification of DWV in the mites showed no significant difference in virus copy numbers between the phoretic mites (V-Pho1) and in the mites reared for two days in the PBS-injected pupae in Experiment 1 (V0-PBS_1_), while two days feeding on pupae injected with DWV (V0-DWV_1_) resulted in a statistically significant 357-fold increase (Fig. 3a, df = 31, F value = 46.63, P < 0.0001, large effect size, Cohen’s *d* =2.40). Importantly, in both Experiments 1 and 2, significantly higher DWV levels were observed in the mites fed on the DWV-injected pupae with high DWV levels, (V0-DWV_1_ and V0-DWV_2_) compared to their respective control mites fed on PBS-injected pupae with significantly lower DWV levels (V0-PBS_1_ and V0-PBS_2_), a 625 fold increase in Experiment 1 (Fig. 3a, df = 39, F value = 51.55, P < 0.0001; large effect size, Cohen’s *d* = 2.27), and a 51-fold increase in Experiment 2 (Fig. 4a; df = 23, F value = 12.66, P = 0.001760, large effect size, Cohen’s *d* =1.45).

Surprisingly, high DWV loads in the mites (V0-DWV_1_ and V0-DWV_2_) fed for 2-days on pupae with high DWV levels (P0-DWV_1_ and P0-DWV_2_), were not maintained for the duration of the experiment. In both experiments the levels of DWV in these mites decreased significantly after four 2-day passages on un-injected pupae with low DWV levels, with a 29-fold decrease in Experiment 1 (V0-DWV_1_ and V4-DWV_1_; Fig, 3a, df = 40, F value = 32.19, P < 0.0001; large effect size, Cohen’s *d* = 1.77), and a 18-fold decrease in Experiment 2 (V0-DWV_2_ and V4-DWV_2_; Fig. 4a; df = 21, F value = 28.65, P < 0.0001; large effect size, Cohen’s *d* =2.28). After the passages, the levels of DWV in the mites initially exposed to the high DWV-level pupae (V4-DWV_1_ and V4-DWV_2_) were not significantly different from those in mites initially reared on control PBS-injected pupae (V4-PBS_1_ and V4-PBS_2_; Fig. 3a, Fig. 4a; P>0.05).

### DWV loads in *Varroa* mites strongly influence virus transmission to the honey bees

To investigate how *Varroa* mites’ ability to transmit DWV was influenced by their DWV loads, we designed Experiment 2 (Fig. 2). Here, DWV levels were quantified in pupae incubated for additional 3 days after mite removal to allow development of the DWV infection. We tested *Varroa*-mediated transmission in the Pupae 1 and Pupae 4 groups, which were exposed to the mites fed on the Pupae 0 group injected with PBS (P1-PBS-Ext_2_ and P4-PBS-Ext_2_) or with DWV (P1-DWV-Ext_2_ and P4-DWV-Ext_2_) (Fig. 2). We observed significantly higher DWV copy numbers in pupae exposed to mites that had directly fed on high DWV-level pupae (P1-DWV-Ext_2_) compared to pupae exposed to mites subjected to four 2-day passages (33-fold difference) after initial rearing on the DWV-infected bees (P4-DWV-Ext_2_) (Fig. 4c, df = 21, F value = 14.94, P = 0.000964; large effect size, Cohen’s *d* =1.79). Importantly, there were no significant differences between DWV levels in pupae from group P4-DWV-Ext_2_, and in the control groups P1-PBS-Ext_2_ and P4-PBS-Ext_2_ (Fig. 4c). A significantly higher proportion of honey bees in the group P1-DWV-Ext_2_ pupae developed a high-level DWV infection above 10^9^ DWV genome copies per bee (7 out of 12), compared to pupal group P4-DWV-Ext_2_ subjected to passaged mites where only 1 pupa out of 10 just met the minimum threshold (Fig. 4c; contingency table analysis, Fisher exact probability test, P=0.0263). Notably, high DWV levels were not observed in pupae exposed to control mites originally fed on PBS injected pupae (P1-PBS-Ext_2_ and P4-PBS-Ext_2_; Fig. 4c).

The use of a clone-derived DWV isolate with known sequence helped trace the source of DWV transmitted by the *Varroa* mites. Sequencing 1.2 kb-long RT-PCR fragments of the DWV genome covering the most divergent leader protein-coding region, amplified from the pooled P0-DWV_2_ and P1-DWV_2_ pupae, showed that DWV in the both pupal groups was identical to the DWV-304 used for the inoculation of the Pupae 0 (Supplementary Data Fig. S1). Sanger sequencing runs showed no polymorphism in both P0-DWV_2_ and P1-DWV_2_ (Supplementary Data Fig. S2), clearly indicating that only the DWV-304 isolate used for infection of the P0-DWV_2_ pupae was transmitted. Contrary to this, the DWV RT-PCR fragment amplified from the V0-DWV_2_ mites was highly polymorphic (Supplementary Data Fig. S2). Due to this, it was not possible to obtain a complete sequence of this fragment, but the terminal sections were also identical to the DWV-304 (Supplementary Data Fig. S1).

### Negative-strand DWV RNA detected exclusively in *Varroa* mites fed on high DWV-level pupae

Quantification of negative-strand DWV RNA was carried out in honey bee and mite samples pooled according to their treatment (in this analysis we excluded a single mite exposed to pupae with high DWV levels from the pool V0-PBS_1_, Fig. 3a) using a tagged primer approach. Two independent assays targeting different regions in the DWV genome and using different primer tags were used (Supplementary Data Table S1). Accuracy of these tests was confirmed by detection of negative-strand DWV RNA in the honey bee pupae pools containing individuals with more than 10^9^ DWV genome copies (Fig. 4d). In both assays, which included melting curve analysis to confirm the identities of amplified fragments, the negative-strand DWV RNA was detected exclusively in the mites directly fed on the pupae with high DWV levels for 2 days. This included V0-DWV_1_ mites from Experiment 1 (Fig. 3b) and V0-PBS_2_ and V0-DWV_2_ mites from Experiment 2 (Fig. 4b). No negative-sense DWV RNA was detected in mites after four passages on low DWV level pupae in both Experiments 1 and 2 or in phoretic mites, V-Pho1 (Figs. 3b and 4b).

### Detection of honey bee mRNA in *Varroa* mites suggests acquisition of honeybee cells

Accumulation of negative-strand DWV RNA in mites could be explained by the acquisition of honey bee cells, containing the replication complexes^27^ including negative-strand DWV RNA, rather than by replication of DWV in the mite itself. It is possible that during their feeding on the honey bees, *Varroa* mites consume not only pure, “filtered” hemolymph plasma, but also honey bee cells. Further, the digestion of these honey bee cells in the *Varroa* midgut could explain the disappearance of the negative-strand DWV RNA following passage on low DWV-level pupae. Indeed, the detection of honey bee transcripts beta-actin (BeeBase gene GB44311) and Apolipophorin-III-like (BeeBase gene GB55452) in the *Varroa* mites using RT-qPCR supports this explanation (Supplementary Data Table 2). We further detected numerous honey bee transcripts in *Varroa* RNA-seq libraries produced in our laboratory (Supplementary Data Table 3), and a different laboratory^28^ (Supplementary Data Table 4). We found that in the *Varroa* RNA-seq libraries the numbers of honey bee beta-actin mRNA were as high as 3.01% ± 0.94% (average ± SD) of those of *Varroa* beta-actin mRNA (Supplementary Data Table 3). There was variation in the proportions of honey bee transcripts present in the mites reared on different honey bee life stages (pupae or adult bees), which might reflect differences in relative expression of these genes in the source honey bee pupae. We also observed higher (about 8-fold) relative levels of total honey bee mRNA transcripts in *Varroa* mites after 15 hours of feeding on the honey bee pupae or adults compared to the mites which were starved for 12 hours (Supplementary Data Table 3). These observed dynamics of honey bee mRNA levels in mites could be explained by a gradual reduction of the ingested honey bee cells. Single-stranded mRNA transcripts are unlikely to be present in hemolymph plasma, therefore their presence likely indicates ingestion of the honey bee cells in addition to intake of “filtered” hemolymph plasma. Indeed, by using RT-qPCR we demonstrated that following freeze-thaw disruption of hemolymph cells the level of beta-actin mRNA decreased about 1000-fold in 2 hours (Supplementary Data Fig. S3) suggesting it is quickly degraded outside the cell.

## Discussion

A growing body of evidence suggests that spread of the mite *Varroa destructor* has greatly increased pathogenicity of DWV, elevating it to the most important viral pathogen of the Western honey bee, *Apis mellifera*^24^. Knowledge of the interaction between DWV and its mite vector is required to better understand the impact of mite vectoring on virus evolution, development of accurate predictive models of *Varroa* mite and DWV dynamics, and design of novel DWV control approaches. Surprisingly, despite the importance of DWV-*Varroa* interactions, it is not clear if replication of all DWV types occur in *Varroa* mites^17^. The main argument in support of the replication of DWV in *Varroa* mites remains the detection of negative-strand DWV RNA in *Varroa* mites, in particular, in mites associated with the pupae with high DWV levels^18, 19^.

Here we used a recently developed *in vitro* system for maintaining *Varroa* mites on honey bee pupae^24^ and a cloned isolate of DWV type A (E. Ryabov, personal communication) to experimentally establish whether DWV replication occurs in *Varroa* mites, to determine factors affecting DWV levels in the *Varroa* mites, and to assess the impact on DWV levels on the mites’ potential for DWV vectoring.

We demonstrated that DWV loads in mites are highly dynamic and rapidly increase or decrease depending on the levels of DWV in the honey bees on which they are feeding on. For example, we used phoretic mites to show that DWV loads increased over 50-fold following 2 day of feeding on pupae pre-infected with DWV, while DWV levels remained unchanged in mites fed on the low-DWV PBS-injected pupae (Fig. 3a, Fig. 4a). The phoretic mites used in this experiment (V-Pho_1_), had relatively low DWV levels (Fig. 3a) despite being collected from DWV-infected colonies, which could be explained by loss of DWV loads following passage on the adult bees, mostly nurses, which usually have low DWV levels. We observed a strong positive correlation between levels of DWV in pupae and their associated *Varroa* mites (Pearson R^2^=0.58; Fig. 5), suggesting that DWV loads in the mites could be explained by virus acquisition from the pupae during feeding. This observation agrees with a previous study which also found a positive correlation between DWV levels in honey bee pupae and their associated mites sourced directly from capped cells from an intact colony^8^. Similar positive correlations between the DWV levels in bees and associated mites observed during natural infestation, where mites were infecting the pupae and were feeding for 5 to 8 days, and in experiments where pupae were pre-infected with DWV and feeding continued only for two days (Figs. 1a and 2) suggest that most of the DWV present in mites in the course of natural infestation is also acquired from the pupae. Importantly, DWV loads in the mites reared on DWV-infected pupae were significantly reduced 32- and 18-fold in Experiment 1 and 2, respectively, following four 2-day passages on low DWV-level pupae (Figs. 3a and 4a). Moreover, after the passages, no significant differences were observed between the DWV loads in mites which previously acquired high DWV doses and those initially exposed to the control PBS-injected pupae (Fig. 3a, Fig. 4a).

**Figure 5.**
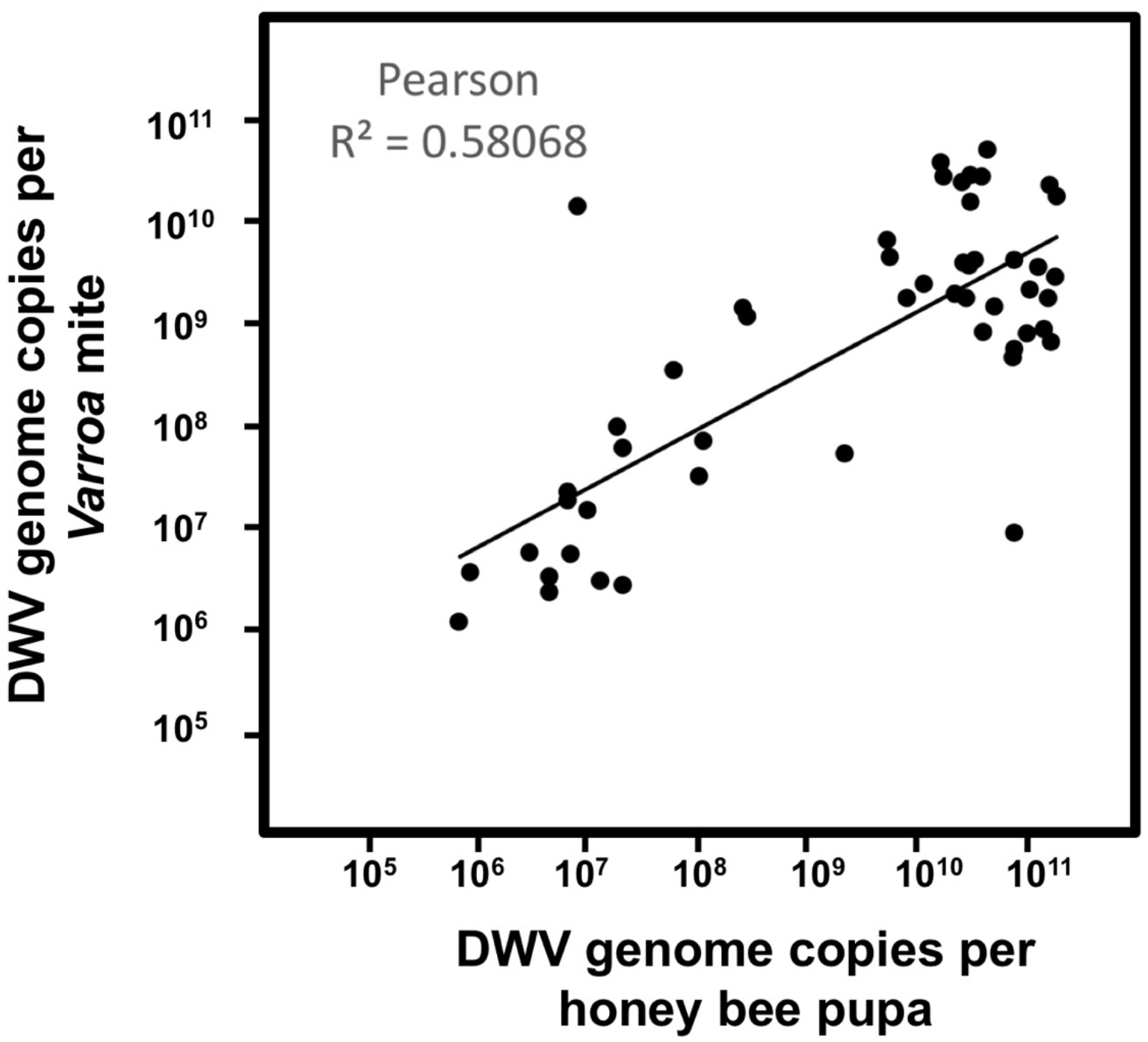
Prevalence of DWV in honey bees (X axis) and associated *Varroa* mites (Y-axis) in simultaneously sampled pupae and mites. Positive correlation between DWV copy numbers (Pearson R^2^ = 0.58068, n=51, p<0.0001). Only mites and pupae with DWV levels above the detection thresholds were used.

Experiment 2 (Figs. 2 and 4c) clearly demonstrated transmission of DWV acquired by mites during feeding on high-DWV level pupae P0-DWV_2_ to the recipient pupae P1-DWV-Ext_2_ by using a cloned DWV isolate. This isolate made it possible to demonstrate mite transmission of the strain that replicated to high levels in the pupae P0-DWV_2_ (Supplementary Data Fig. S1). We further showed that the ability of the mites to vector DWV decreases following passages on low DWV pupae. Significantly lower levels of DWV were observed in the P4-DWV-Ext_2_ pupae fed on by the passaged V4-DWV_2_ mites compared to P1-DWV-Ext_2_ pupae fed on by mites transferred immediately from highly infected pupae (Fig. 4c).

Detection of negative-strand RNA of picorna-like viruses, including DWV, is a good indicator of a virus replication. It is considered the strongest evidence in support of replication of DWV in *Varroa* mites^18,19^. Therefore, we sought to detect negative-strand DWV RNA in the mites from this study. By using two independent RT-qPCR assays for negative-strand DWV RNA quantification, we showed its accumulation exclusively in mites sampled immediately following exposure to pupae with high DWV levels (Fig. 3b, and Fig. 4b). Notably, negative-strand DWV RNA was not detectable in the in the phoretic V-Pho1 mites before exposure to the DWV-level pupae or in the P4-DWV_1_ and P4-DWV_2_ mites with decreased overall DWV loads following passage of low DWV-level pupae (Fig, 3b and Fig. 4b). We hypothesized that such rapid emergence and disappearance of negative-strand DWV RNA in the mites is explained not by replication of DWV in *Varroa* cells, but by acquisition of honey bee cells with containing DWV replication complexes with negative-strand viral RNA^27^. Indeed, acquisition of cells is consistent with detection of honey bee genomic DNA in *Varroa* samples during genome sequencing^30^ and has been recently observed during extensive feeding analyses^31^.

To estimate the extent of honey bee cell acquisition by a mite during feeding, we measured honey bee mRNA levels in the mites. Detection of honey bee mRNA transcripts is an indicator of intact cells acquisition by mites because we demonstrated using RT-qPCR that levels of mRNA in the honey bee hemolymph decline by about 1000 times in 2 hours following cell disruption (Supplementary Data Fig. S3). Therefore, the presence of honey bee mRNA revealed by RT-qPCR in the mites of our experiments (Supplementary Data Table S2) and by analyzing RNA-seq libraries (Supplementary Data Tables S3 and S4) strongly suggests acquisition of entire honey bee cells by *Varroa* mites. The phenomenon of hosts undamaged cells and tissues being accumulated within the midgut lumen of a blood sucking parasite has been observed in other members from the superorder *Parasitiformes*, which includes mites and ticks. In the black-legged tick *Ixodes scapularis* and the American dog tick *Dermacentor variabilis* intact erythrocytes were detected up to 6 days post blood meal. It is speculated that for the subclass *Acari*, the midgut is the primary nutrient storage site where ingested cells are kept intact^32^. Our honey bee cell acquisition model could explain emergence and disappearance of negative-strand DWV RNA in the mites in the course of the Experiments 1 and 2 (Figs. 3b, 4b). Indeed, the lack of replication of DWV-A in *Varroa* mites observed in our study is further supported by a previously published immunolocalization study showing detection of DWV antigen exclusively in the mites midgut lumen^22^.

Our finding that replication of DWV, or at least its most common type DWV-A, does not take place in *Varroa* mites provides a novel insight into the impact of the *Varroa* on virus evolution. The main implication is that, in the absence of replication no generation of additional genetic DWV diversity occurs in *Varroa* mites and no adaptation of DWV for replication in mites takes place. Therefore the reported effect of *Varroa*-mediated transmission on DWV genetics^15–17^ is more likely a consequence of selection of isolates with better non-propagative transmission ability, e.g. those with higher stability in mites and/or viral variants better adapted to transmission *via* injection into the hemolymph. Indeed, previous studies have shown that selection of DWV strains could take place without V*arroa* mites, solely by artificial injection into hemolymph^16^. A lack of replication of DWV in *Varroa* mites is also consistent with the absence of the reports a negative impact of DWV on *Varroa* mites, which could be a result if replication of DWV took place in mites.

In light of finding no evidence of DWV persistence in *Varroa* mites, it is possible that development of severe *Varroa* and DWV-associated honey bee colony decline requires colonies to reach a certain threshold level of both *Varroa* mite density and proportion of bees with high DWV levels. Indeed, increased DWV levels were reported in colonies with high *Varroa* mite infestation^13^. Also, the lack of replication and low persistence of DWV in *Varroa* mites could explain the absence of *Varroa* related damage in some island honey bee populations^33^.

It should be highlighted that the reported experiments were carried out using one type of DWV-like viruses, namely DWV type A. This cloned DWV variant^24^ was isolated from a heavily Varroa-infested colony typical for the USA and likely to be Varroa-selected, as evidenced by its transmission in Experiment 2. It cannot be ruled out that other DWV-like variants, namely the highly pathogenic DWV type B^6^, which has shown increased virulence^34,35^, or VDV1-DWV recombinants^8–10^ could have replicative ability in *Varroa* mites, especially considering their preferable transmission by the mites^8,36^ and the detection of VDV-1 negative strand RNA in the *Varroa* synganglions^23^. Investigation of the abilities of different DWV strains to replicate in the mites should be carried out using experimental procedures similar to those used in this study.

Taking into account the implications of a lack of replication of DWV in the *Varroa* mites, it might be possible to control DWV solely by targeting DWV in its bee host (e.g. using an RNAi approach). Data on the rate of reduction of the mites’ ability to transmit DWV following passages on low DWV bees (as it likely to occur in the case of the phoretic mites) would allow for the development of precise predictive model linking *Varroa* infestation rate and DWV load. This would be important for planning measured treatments against *Varroa* using acaricides, which may have negative health impacts on bees^37^.

Results of this study clearly suggest that DWV, or at least the most common DWV type A, is transmitted between honey bees by the *Varroa* mite in a non-propagative manner. Significant declines in the mites’ ability to transmit DWV suggest non-persistent transmission mechanism. While this does not diminish the role played by mites in DWV infection of honey bees, it does clarify the selective forces acting on DWV. Namely, this virus has evolved to exploit the honey bee, not adapted to replication in mite cells, and has a more intimate relationship with honey bee immune defenses than those of the mite. These insights are critical in reducing the impacts of the tripartite relationship among honey bees, mites, and DWV.

## Methods

### *Honey bees and Varroa* mites

Honey bee pupae were obtained from the Beltsville USDA Bee Research Laboratory (BRL) apiary from strong colonies “132”and “Nuc-L” (two boxes with more than 10 frames of brood and stored food) with a low (less than 1%) *Varroa* mite infestation rate, and the DWV loads in the pupae undetectable by qRT-PCR test^17,38^ in June 2018. The colonies were not subjected to any *Varroa* control treatments. The colonies “132”and “Nuc-L” were used in Experiment 1 (July 2018) and in Experiment 2 (September 2018) respectively. Pupa at the white-or pink-eyed stage were pulled out of cells using soft tweezers no more than 24 hour prior their use in the experiments. *Varroa* mites were collected from the same apiary, using colonies showing about 7% *Varroa* mite brood infestation rate and detectable levels of DWV in brood^40^. Phoretic mites collected from adult bees of colonies “3” were used for Experiment 1, colonies “71” and “73” were used as a *Varroa* source in Experiment 2, where both phoretic and founder *Varroa* mites were collected. All *Varroa* mite source colonies used in this study harbored only DWV-A and were free of VDV-1 (DWV-B). The tests for the presence of DWV-A and DWV-B were carried out as described previously^17,38^. The sugar roll method was used to collect *Varroa* mites; this included shaking powdered sugar-dusted adult bees over a tray with water, collecting the released mites from the water with a strainer, drying them on tissue paper, then placing the mites on honey bee pupae within 2 to 3 hours after collection^25^.

The experiments were organized with a completely randomized design using two treatments. Pupae were either injected as previously described^16^ with DWV (cDNA clone-derived DWV-A variant DWV-304^24^, GenBank Accession Number MG831200) or PBS as a control. The virus was propagated in honey bee pupae injected with infectious DWV *in vitro* RNA transcript produced from the linearized template, plasmid DWV-304, using T7 RNA polymerase (HiScribe T7 kit, New England Biolabs) according to manufacturer’s instructions (E. Ryabov, personal communication). For preparation of the clonal DWV extracts containing infectious DWV virus particles, individual transcript-injected pupae were homogenized with 2 mL of phosphate-buffered saline (PBS at 3 days post injection). For each individual pupal extract, 1 mL was used for preparation of the DWV inocula, which included clarification by centrifugation at 3000g for 5 min and filtration through 0.22 μm nylon filter (Millipore). The DWV concentration in the extract was quantified by qRT-PCR and the infectivity of the virus was confirmed by development of typical overt DWV infection, about 10^11^ genome copies per bee. The identity of clonal DWV in the extract with its respective parental DWV cDNA clone was confirmed by sequencing of the RT-PCR fragment corresponding to the leader protein gene of DWV-A (nucleotide positions 950-2050 of the genomic RNA). The DWV extract was stored at −80°C prior to use. Twenty or twelve replicates were used for each treatment in Experiments 1 and 2, respectively. Each replicate consisted of one pupa and two or three *Varroa* mites contained inside a gelatin capsule (Veg K-Caps size 00 capsule from Capsuline, Pompano Beach, FL, USA). *Varroa* mites were tested with a paint brush for mobility to be sure that they were alive. The gelatin capsule with the pupa and mites were inserted into a 32/8-Place Combo Tube Rack (USA Scientific, Orlando. FL, USA) inside an environmental chamber set to 32°C and about 85% of relative humidity in complete darkness. The mites and pupae were checked daily for ten days and survival was recorded. Only live mites were transferred to the next pupae and mite feeding was confirmed by accumulation of their feces in the gelatin capsules. Mite survival was analyzed as previously^25^ using JMP Software (SAS, Cary, NC, USA).

### Quantification of RNA by RT-qPCR

Total RNA was extracted from individual honey bee pupae using Trizol reagent (Ambion) and further purified using a RNeasy kit (QIAGEN) according to manufactures’ instructions. Extraction of total RNA from individual *Varroa* mites was carried out using a RNeasy kit (QIAGEN) and included disruption of the mites in the RLT buffer supplied with the RNeasy kit using a glass micro-sized 0.2 mL tissue grinder (DWK Life Sciences, Wheaton), followed by further disruption of the mite tissues using QIAShredder (QIAGEN). DWV RNA loads in individual honey bee pupae and associated *Varroa* mites were quantified by reverse transcription quantitative PCR (RT-qPCR), which included cDNA synthesis using random hexanucleotides. Detection mainly included the genomic positive strand DWV RNA (Supplementary Data Table S1). RT-qPCR was carried out essentially as previously described^28,39^, DWV RNA copy numbers were log-transformed prior to statistical analyses. Negative-strand DWV RNA quantification was carried using two sets of primers designed for targeting different regions in the DWV genome, Assay 1 (region 4909-5008 nt)^16^ and Assay 2 (region 7303-7503 nt)^40^.

### mRNA degradation assay

Hemolymph cell suspension samples were collected from 10 pink eye honey bee pupae, approximately 30 μL in total. A 5 μL sample was used to extract total RNA immediately following collection. The remainder of the sample was subjected to 5 cycles of freezing and thawing (5 minutes each) and vortexing to disrupt cellular membranes and incubated at +33°C. Subsequent 5 μL aliquots, were collected for immediate RNA extraction after 30 min, 60 min, and 90 min of incubation. The extracted RNA samples were used to quantify honey bee beta actin mRNA by RT-qPCR using specific primers (Supplementary Data Table 1).

### Analysis of RNA-seq data

Pre-analysis processing of the raw *Varroa* RNAseq Illumina data (paired-end 150 nt reads) was performed as previously described^39^. Cleaned data from each sample library were individually aligned to a reference set containing *A. mellifera* official gene set 3.2 (amel_OGSv3.2)^41^, *V. destructor* actin-like mRNA (XR_002672520.1), *V. destructor* myosin heavy chain mRNA (XM_022800959.1), *V. destructor* vitellogenin 1 mRNA (JQ974976.1) and the genomic sequence of DWV RNA (GU109335.1), using Bowtie2^42^. SAMtools^43^ idxstats was used to identify the numbers of the reads corresponding to the *A. mellifera*, V*. destructor* and DWV genomic RNA sequences. Additional, previously published *Varroa* RNAseq libraries^28^ (NCBI BioProject PRJNA392105), were screened for the presence of honey bee and *Varroa* mRNAs using BLAST^44^.

## Acknowledgements

This research used resources provided by the SCINet project of the USDA - Agricultural Research Service, ARS project number 0500-00093-001-00-D and was supported by the USDA National Institute of Food and Agriculture grant 2017-06481 to EVR, YPC and JDE. USDA is an equal opportunity provider and employer.

## Author Contributions

F.P.-F. and E.V.R., the two corresponding authors, conceived the study and contributed equally to this research. Y.C. supervised honeybee colony monitoring. F.P.-F and E.V.R. carried work with bees and mites. E.V.R. carried out the laboratory molecular work and analyzed virus quantification. M.H. analyzed accumulation of the honey bee transcripts. A.K.C. analyzed the NGS data. F.P.-F. A.K.C., Y. C, J.D.E. and E.V.R., wrote the manuscript. All co-authors contributed to data interpretation, and to the writing of the manuscript.

## Supplementary Information

**Supplementary Table 1.**
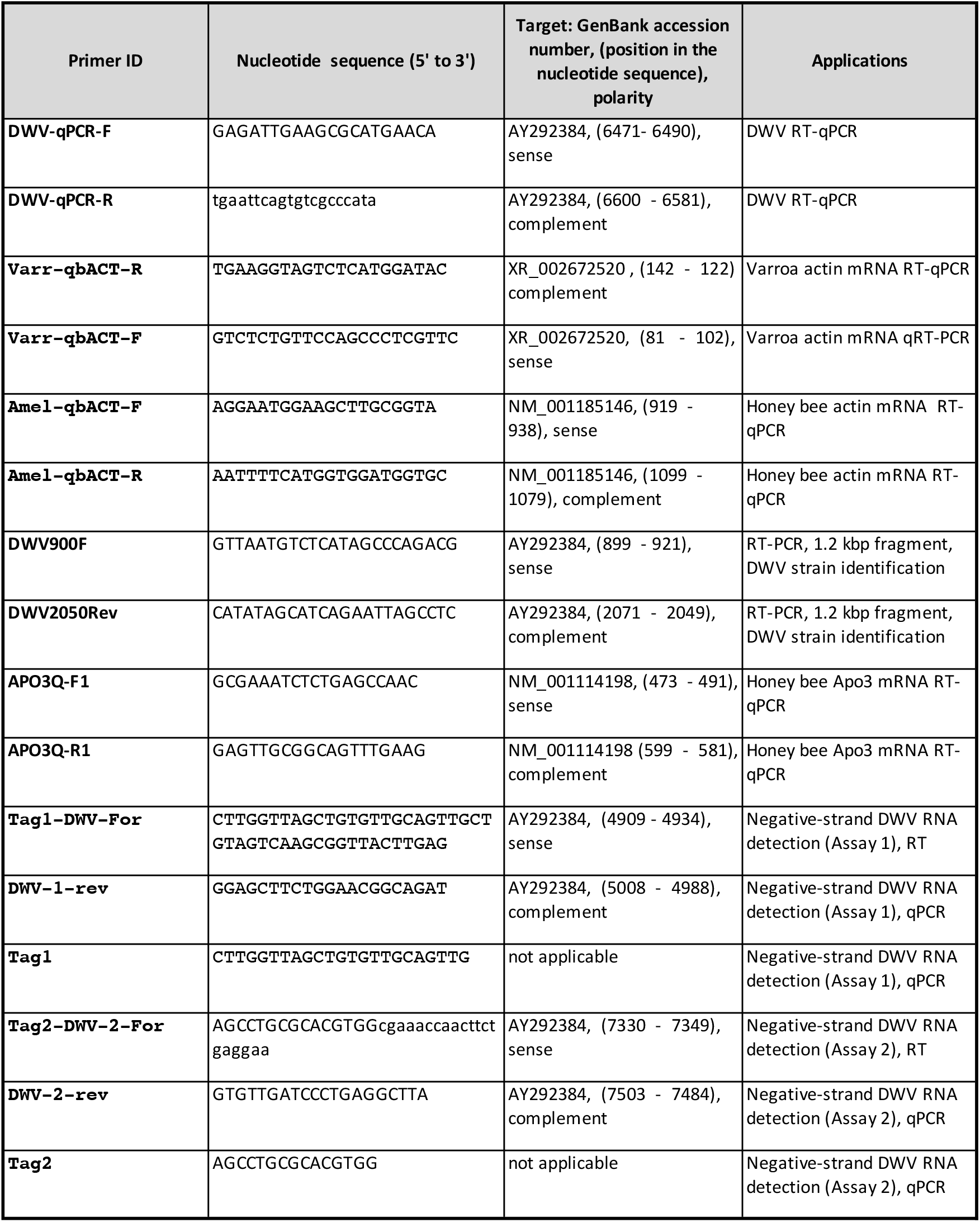
Primers used in this study.

**Supplementary Table 2.**
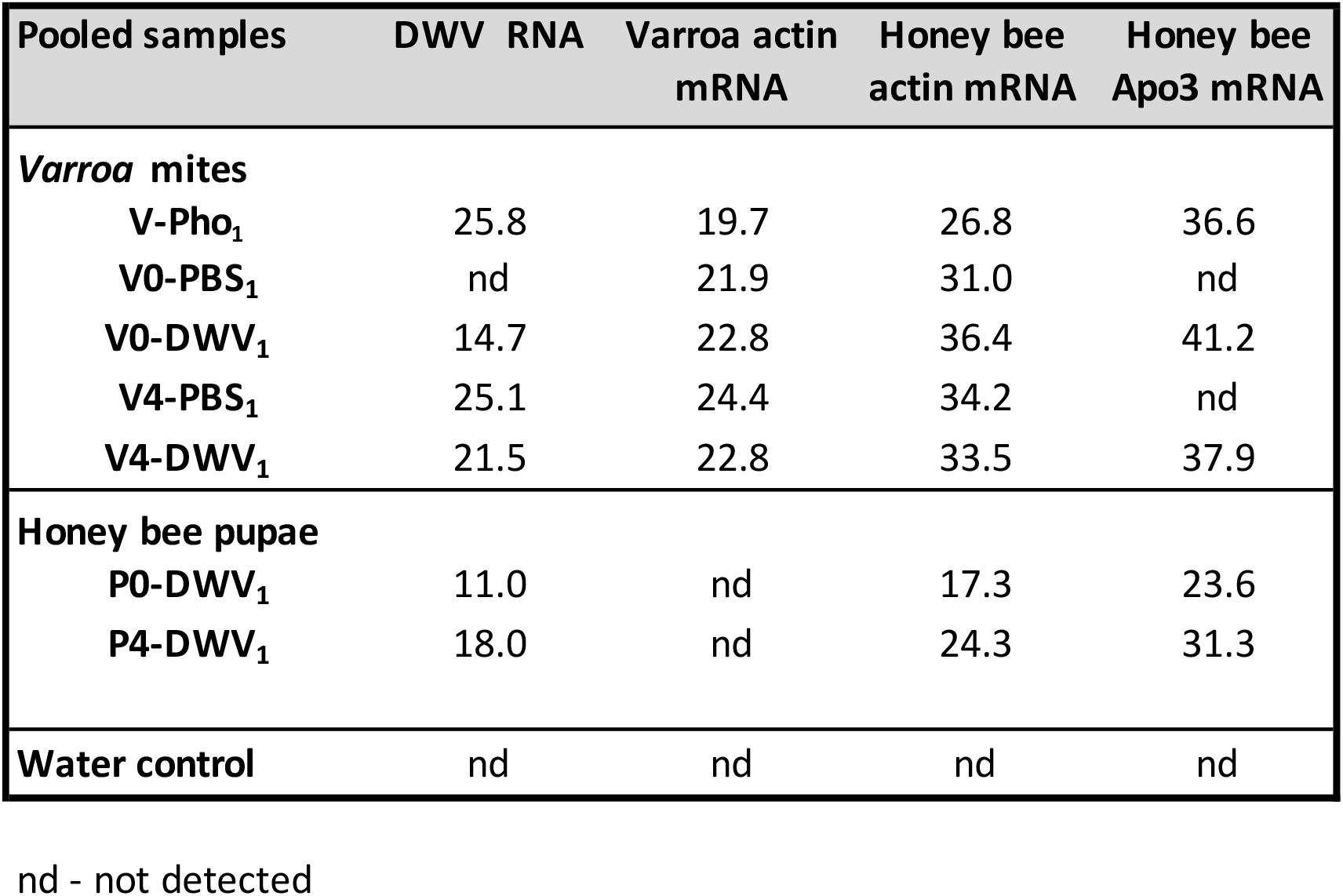
Detection of honey bee mRNA in *Varroa* mites by RT-qPCR. Ct values for honey bee actin (BeeBase gene identifier GB44311) and Apolipophorin-III-like, Apo3 (BeeBase gene identifier GB55452) mRNAs, *Varroa destructor* actin (GenBank Accession XR_002672520) mRNA, and DWV genomic RNA in pooled *Varroa* mite and control honey bee pupae.

| Pooled samples | DWV RNA | Varroa actin mRNA | Honey bee actin mRNA | Honey bee Apo3 mRNA |
| --- | --- | --- | --- | --- |
| ***Varroa* mites** |  |  |  |  |
| V-Pho<sub>1</sub> | 25.8 | 19.7 | 26.8 | 36.6 |
| V0-PBS<sub>1</sub> | nd | 21.9 | 31.0 | nd |
| V0-DWV<sub>1</sub> | 14.7 | 22.8 | 36.4 | 41.2 |
| V4-PBS<sub>1</sub> | 25.1 | 24.4 | 34.2 | nd |
| V4-DWV<sub>1</sub> | 21.5 | 22.8 | 33.5 | 37.9 |
| **Honey bee pupae** |  |  |  |  |
| P0-DWV<sub>1</sub> | 11.0 | nd | 17.3 | 23.6 |
| P4-DWV<sub>1</sub> | 18.0 | nd | 24.3 | 31.3 |
| **Water control** | nd | nd | nd | nd |

nd - not detected

**Supplementary Table 3.** Detection and quantification of honey bee mRNAs in RNAseq libraries from phoretic *Varroa* mites following starvation or feeding. Numbers of NGS reads corresponding a set of *Varroa* and honey bee mRNAs as well as DWV genomic RNA were determined in *Varroa destructor* RNAseq libraries (Egekwu & Cook, submitted).

| Varroa RNA-seq library ID |  | Var_C333_VaD_WaS_12H | Var_C3_VaD_WaF_15H | Var_C3_VaD_WpF_15H |
| --- | --- | --- | --- | --- |
| Varroa mite treatment |  | Phoretic mites starved for 12 hours | Phoretic mites fed on adult honey bees for 15 hours | Phoretic mites fed on honey bee pupae for 15 hours |
| Total number of reads in the library |  | 59560566 | 49673302 | 65165306 |
| DWV reads |  | 38870677 | 18964968 | 24572362 |
| Reads numbers ratio, honey bee beta-actin mRNA to *Varroa* beta-actin |  | 0.017 | 0.039 | 0.034 |
| Reads numbers ratio, total honey bee mRNAs to *Varroa* beta-actin mRNA |  | 2.540 | 24.152 | 19.066 |
| *Apis mellifera* mRNA | *A. mellifera* beta-actin (GB44311) NM_001185146 | 626 | 674 | 836 |
|  | *A. mellifera* myosin heavy chain (GB51653) | 1428 | 4094 | 5231 |
|  | *A. mellifera* apolipoprotein III-like protein (GB55452) | 1 | 240 | 220 |
| Total *A. mellifera* mRNAs reads |  | 92604 | 414034 | 470256 |
| *Varroa destructor* mRNA | XR_002672520, *V. destructor* beta actin | 36462 | 17143 | 24665 |
|  | *V. destructor* myosin heavy chain | 2246 | 3318 | 4859 |
|  | *V. destructor* vitellogenin 1 JQ974976 | 1901 | 773 | 1289 |

**Supplementary Table 4.** Detection of honey bee mRNAs in RNAseq libraries from different *Varroa* mite developmental stages. Numbers of NGS reads corresponding a set of *Varroa* and honey bee mRNAs were determined in *Varroa destructor* RNAseq libraries^23^ (NCBI BioProject PRJNA392105).

| *Varroa destructor* RNA-seq library, accession number, mite description |  | SRX2960636<br>Emerging mites<br>(collected from P8 to P9 brood cells) | SRX29606371<br>Young mites<br>(collected from P8 to P9 brood cells) | SRX2960647<br>Young mites<br>(collected from P8 to P9 brood cells) | SRX2960634<br>Laying mites<br>(collected from sealed brood cells containing pre-pupae) | SRX2960649<br>Phoretic mites<br>(collected on adult bees) | SRX2960670<br>Phoretic mites<br>artificially reared in cages with adult bees | SRX2960669<br>Phoretic mites<br>(collected on adult bees) | SRX2960632<br>Arresting mites<br>(collected in unsealed L5 brood cells) |
| --- | --- | --- | --- | --- | --- | --- | --- | --- | --- |
| Total number of reads in the library |  | 26531852 | 27083570 | 33433690 | 26670037 | 17467280 | 29446920 | 32563456 | 28709594 |
| *Apis mellifera*<br>mRNAs | NM_001185146 *A. mellifera*<br>actin related protein 1 (Arp1) | 140 | 473 | 683 | 174 | 93 | 206 | 88 | 87 |
|  | XM_02644216 *A. mellifera*<br>myosin heavy chain | 122 | 758 | 355 | 36 | 56 | 92 | 57 | 14 |
|  | NM_001114198 *A. mellifera*<br>apolipoprotein III-like protein | 3 | 51 | 77 | 11 | 1 | 0 | 0 | 2 |
| *Varroa destructor*<br>mRNAs | XR_002672520, *V. destructor*<br>actin-like protein | >20000 | >20000 | >20000 | 8606 | 17838 | >20000 | 13931 | 9678 |
|  | XM_022800959 *V. destructor*<br>myosin heavy chain | 2314 | 2523 | 4080 | 1847 | 1494 | 2920 | 1017 | 1061 |
|  | JQ974976 *V. destructor*<br>vitellogenin 1 | 1649 | 524 | 1182 | >20000 | 848 | 1446 | 756 | 3405 |

**Supplementary Figure S1.**
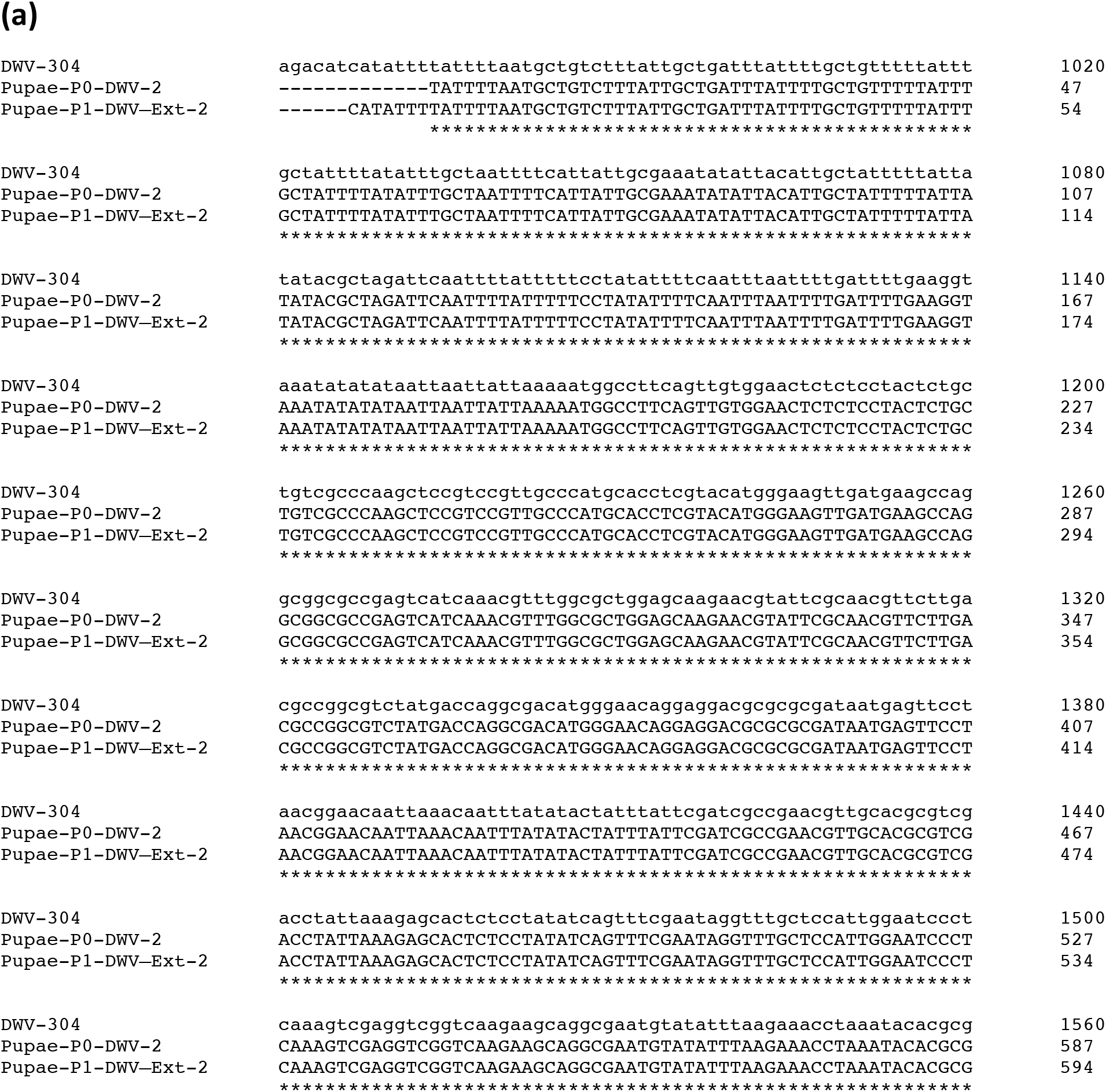
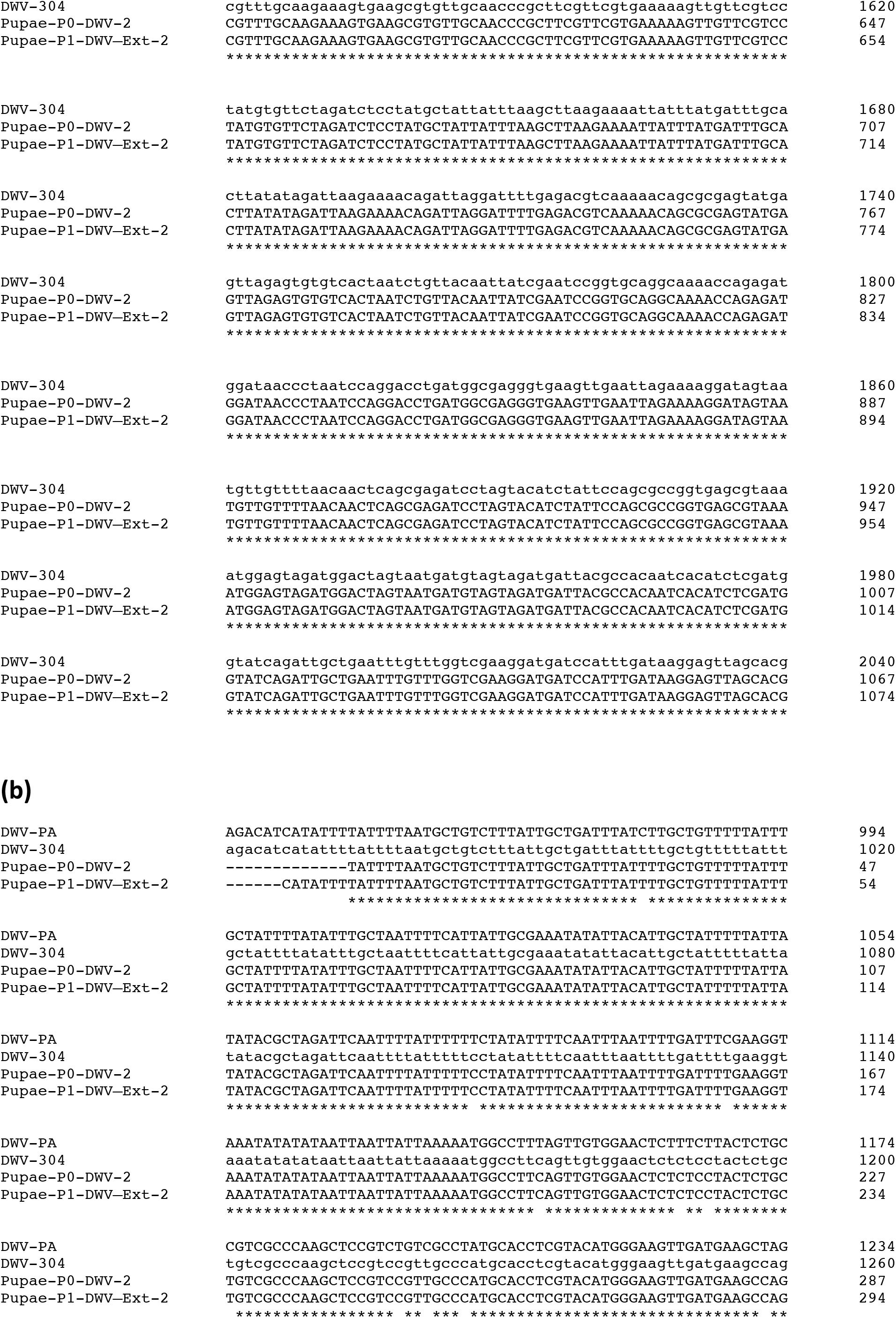
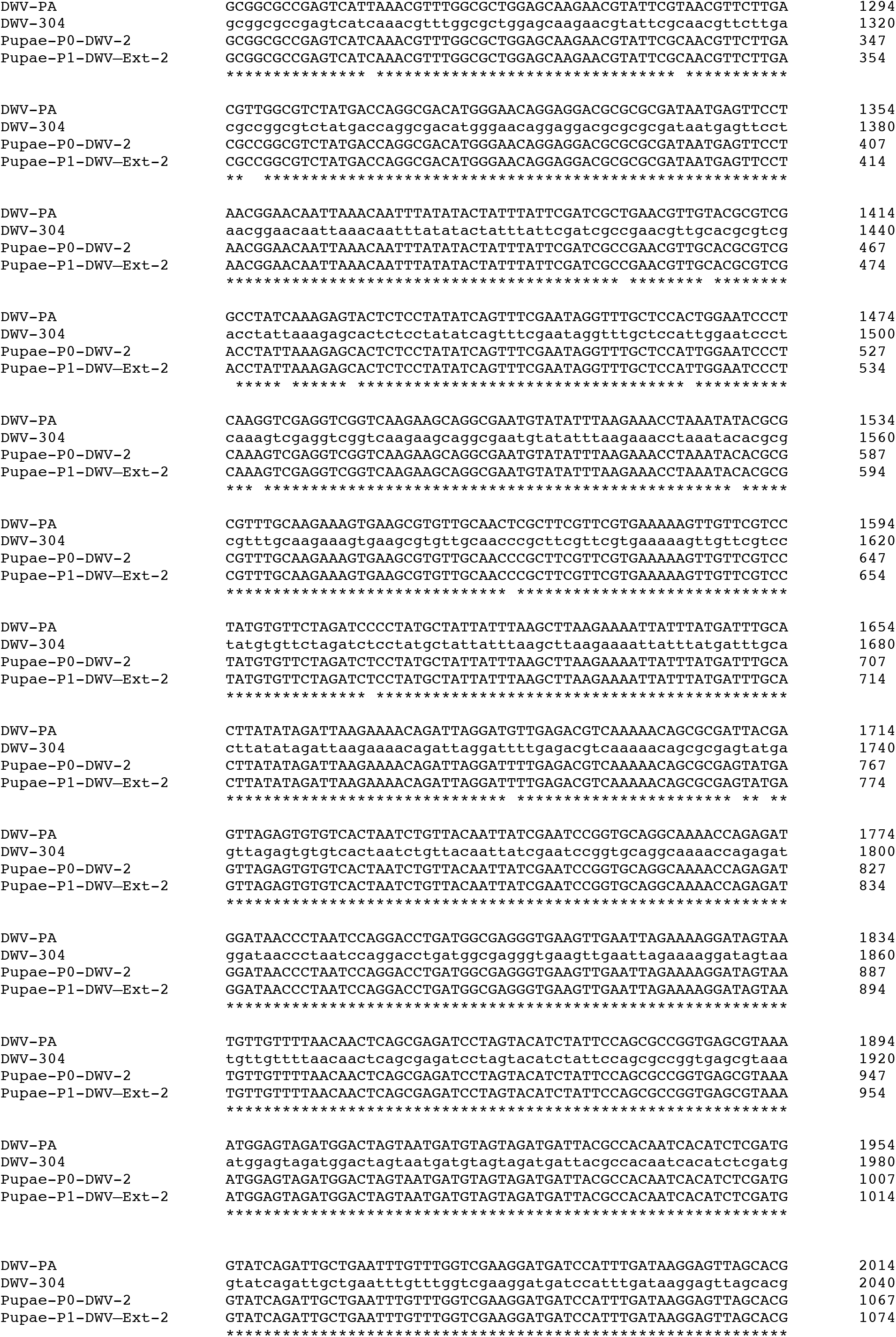
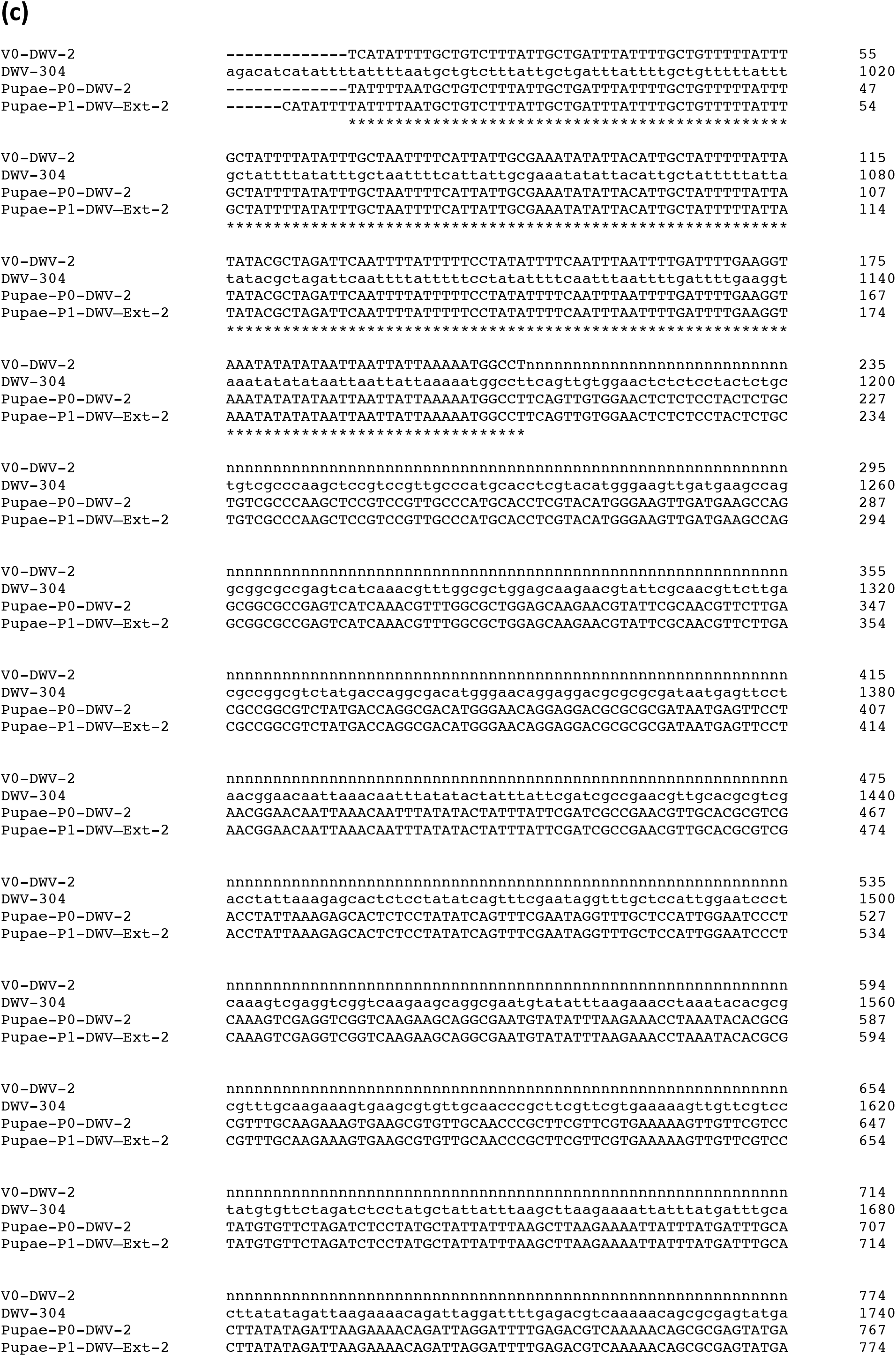
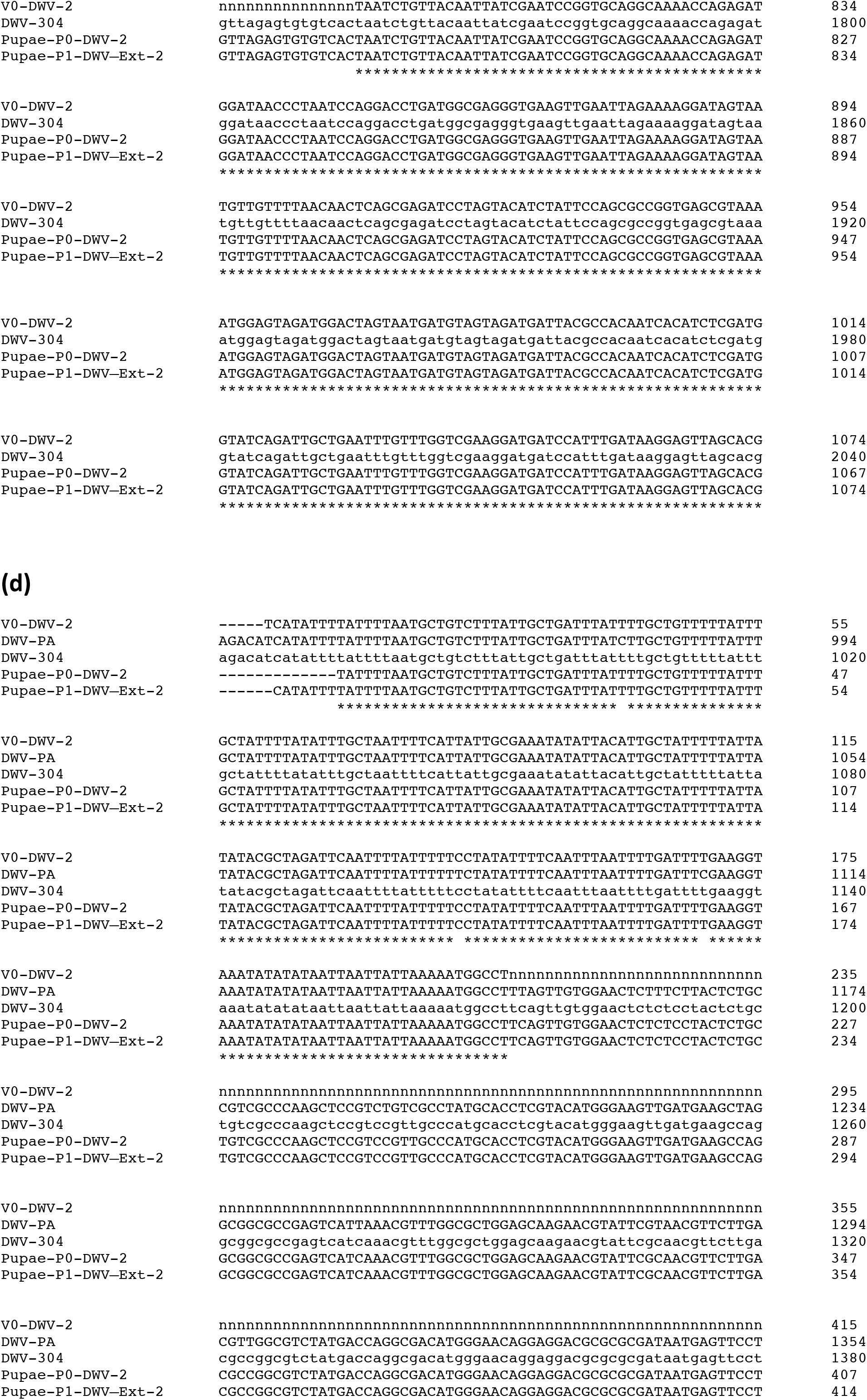
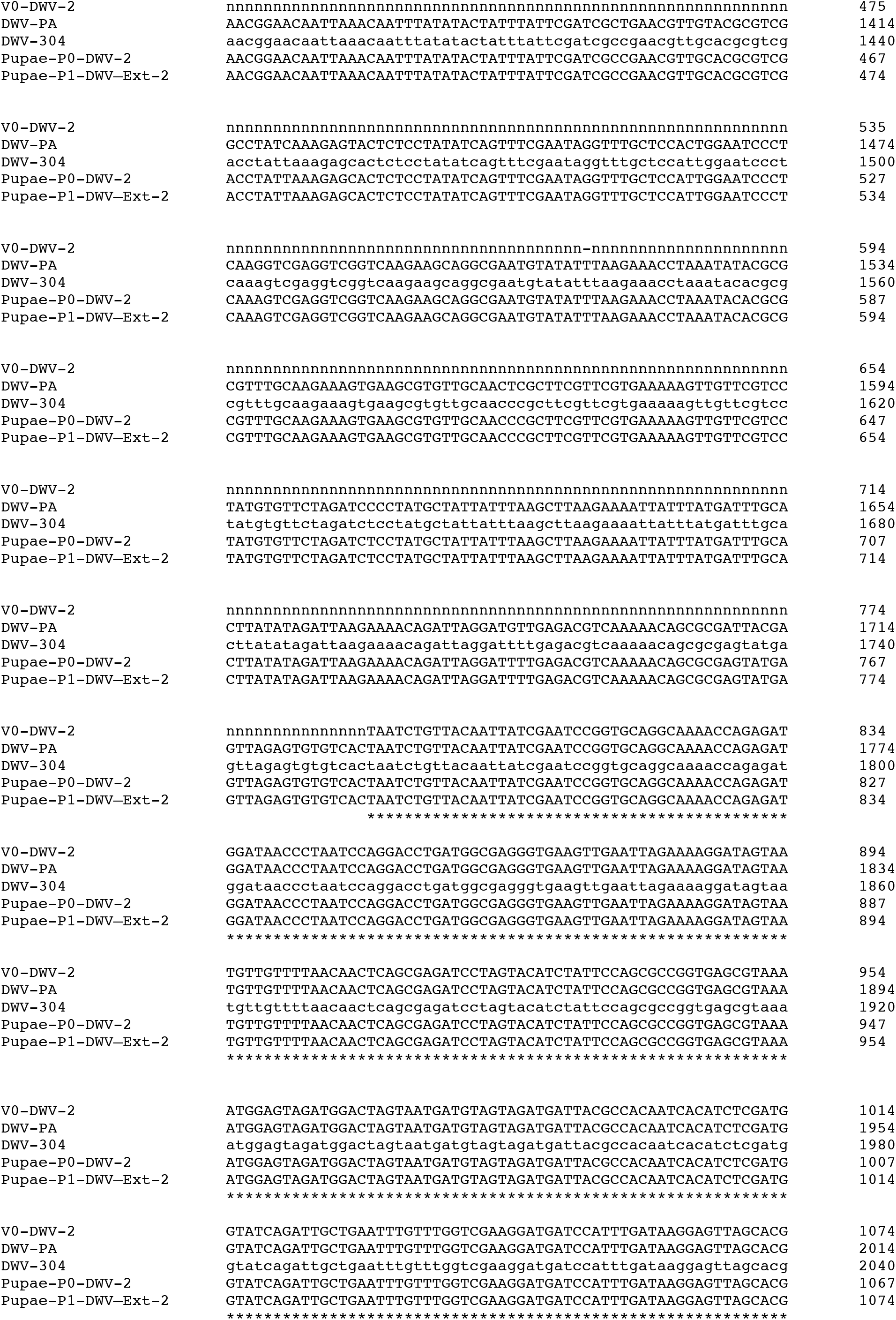
Alignments of consensus DWV RNA nucleotide sequences from honey bees and mites in Experiment 2: “*Varroa-*mediated DWV transmission experiment”. RT-PCR fragments corresponding to the DWV LP region were amplified using RNA extracted from bees and mites pooled according to their treatment (Experiment 2, Figs. 2 and 4). **(a, b)** Complete identity between the DWV sequences amplified from honey bee pupae P0-DWV_2_, P1-DWV_2_, and the cloned isolate DWV-304. **(c, d)** Complete nucleotide identity between the sequenced terminal portions of the RT-PCR fragment amplified from V0-DWV_2_ mites, the sequences from honey bee pupae P0-DWV_2_, P1-DWV_2_, and the cloned isolate DWV-304. Due to high polymorphism in the mite DWV load only the terminal portions of V0-DWV_2_ were sequenced. Cloned DWV isolate DWV-304 (GenBank accession number MG831200, isolated in the USA in 2015) was used for pupae P0-DWV_2_ injection. The type DWV isolate DWV-PA isolated the USA in 2006 (GenBank Accession number AY292384), was included with alignments **(b)** and **(c)** to show distribution of polymorphic sites in the LP region of DWV RNA.

**Supplementary Figure S2.**
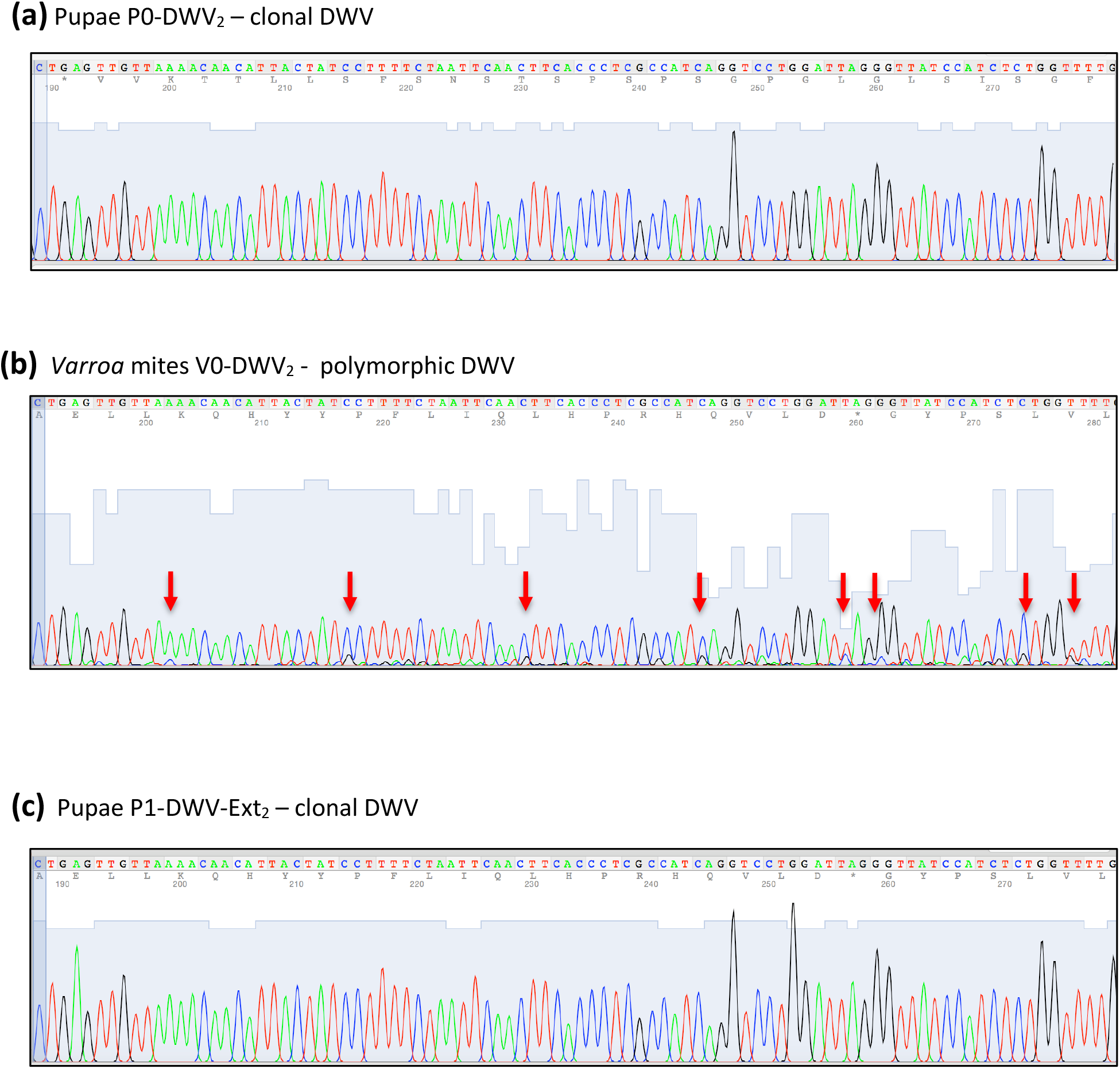
DWV polymorphisms in honey bees and mites in Experiment 2: “*Varroa-*mediated DWV transmission experiment”. Electropherograms of direct Sanger sequencing of RT-PCR fragments corresponding to the DWV LP region were amplified using RNA extracted from bees and mites pooled according to their treatment (Experiment 2, Figs. 2 and 4). **(a**) Honey bee pupae P0-DWV_2_, infected with a clone-derived DWV isolate served as the “source bees”; **(b)** *Varroa* mites V0-DWV_2_, which acquired DWV from the “source bees”; **(c)** Honey bee pupae P1-DWV-Ext_2_ infected with DWV strains transmitted by the *Varroa* mites fed first on the “source bees”. Clonal DWV accumulated in the honey bee pupae and divergent DWV accumulated in the *Varroa* mites, as evidenced by the presence of double peaks indicated by arrows.

**Supplementary Figure S3.**
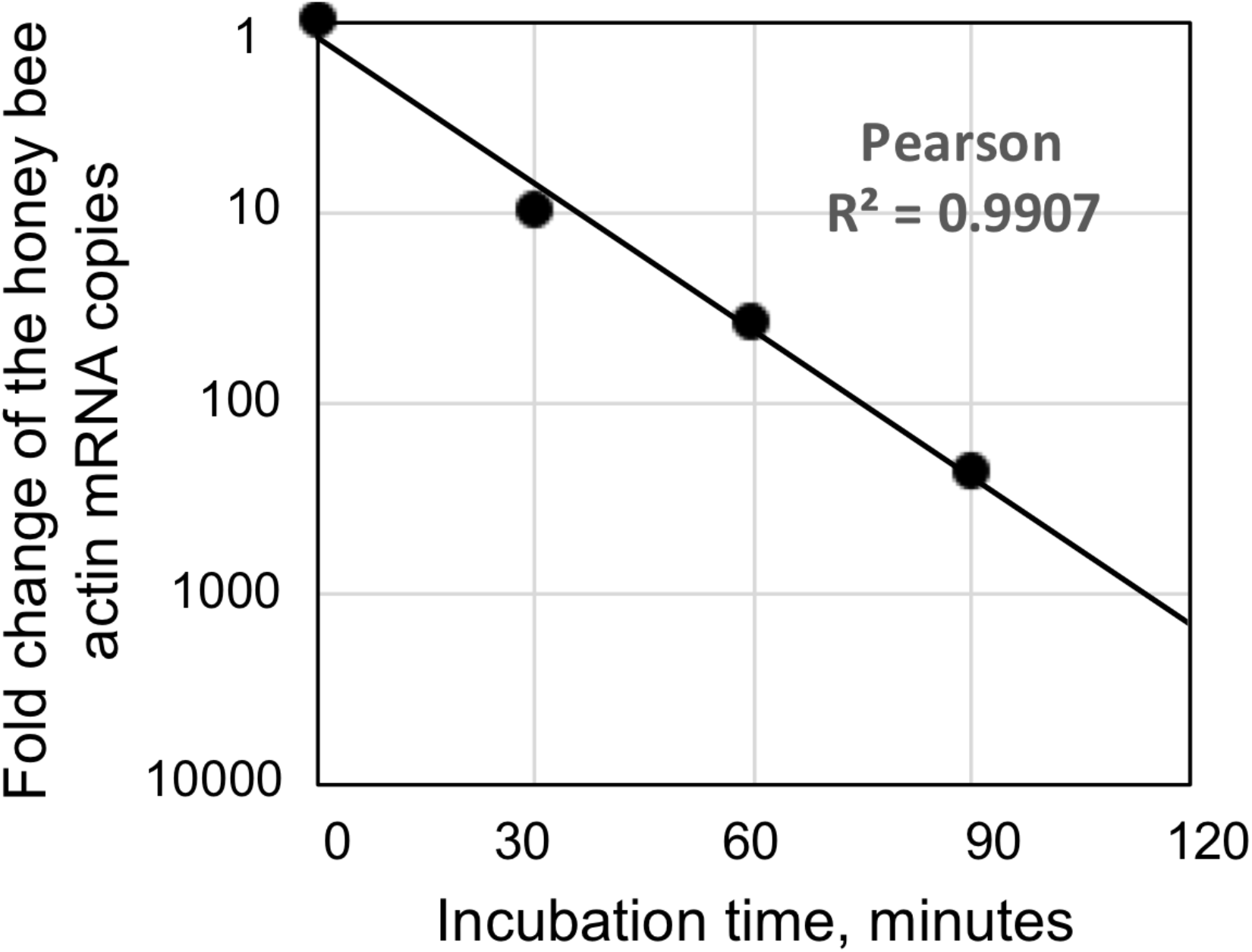
Single stranded RNA degradation in disrupted honey bee hemolymph cells.

